# *KALRN* Mutations Promote Anti-tumor Immunity and Immunotherapy Response in Cancer

**DOI:** 10.1101/2020.01.28.922682

**Authors:** Mengyuan Li, Yuxiang Ma, You Zhong, Lei Qiang, Xiaosheng Wang

**Author notes:** Correspondence to: Xiaosheng Wang (E-mail address). Equal contribution (Mengyuan Li, Yuxiang Ma).

## Abstract

**Background:** *KALRN* (kalirin RhoGEF kinase) is mutated in a wide range of cancers. Nevertheless, the association between *KALRN* mutations and the pathogenesis of cancer remains unexplored. The identification of biomarkers for cancer immunotherapy response is important considering that immunotherapies show beneficial effects only in a subset of cancer patients.

**Methods:** We explored the correlation between *KALRN* mutations and anti-tumor immunity in 10 cancer cohorts from The Cancer Genome Atlas (TCGA) program by the bioinformatics approach. Moreover, we verified the findings from bioinformatics analysis by in vitro experiments. Furthermore, we explored the correlation between *KALRN* mutations and immunotherapy response in four cancer cohorts receiving immune checkpoint blockade therapy.

**Results:** We found that anti-tumor immune signatures were stronger in *KALRN*-mutated than in *KALRN*-wildtype cancers. Moreover, *KALRN* mutations correlated with increased tumor mutation burden and the microsatellite instability or DNA damage repair deficiency genomic properties which may explain the elevated anti-tumor immunity in *KALRN*-mutated cancers. Furthermore, we found that PD-L1 expression was significantly upregulated in *KALRN*-mutated versus *KALRN*-wildtype cancers. The enhanced anti-tumor immune signatures and PD-L1 expression in *KALRN*-mutated cancers may favor the response to immune checkpoint blockade therapy in this cancer subtype, as evidenced in four cancer cohorts receiving anti-PD-1/PD-L1/CTLA-4 immunotherapy. We further revealed that the significant association between *KALRN* mutations and increased anti-tumor immunity was attributed to that *KALRN* mutations compromised the function of KALRN target Rho GTPases on regulating DNA damage repair pathways.

**Conclusions:** The *KALRN* mutation is a useful biomarker for predicting the response to immunotherapy in cancer patients.

## 1. Introduction

Recently, immunotherapy has achieved success in treating various cancers ^1-4^. In particular, the immune checkpoint blockade (ICB), such as inhibition of CTLA-4 (cytotoxic T-lymphocyte-associated protein 4), PD-1 (programmed cell death protein 1), and PD-L1 (programmed cell death 1 ligand), has been used for therapy of many advanced malignancies ^5^. Nevertheless, many cancer patients failed to respond to the immunotherapy. To this end, numerous studies have identified certain genetic or genomic features that are associated with cancer immunotherapy response, such as PD-L1 expression ^6^ and tumor mutation burden (TMB) ^7^. The mutations of certain genes, such as *TP53* ^8^ and *ARID1A* ^9^, have been associated with cancer immunotherapy response.

*KALRN* (kalirin RhoGEF kinase) encodes a protein which activates specific Rho GTPase family members to regulate neurons and actin cytoskeleton ^10^. This gene are mutated in a wide type of cancers ^11^, e.g., melanoma, lung cancer, uterine corpus endometrial carcinoma (UCEC), glioblastoma multiforme (GBM), and colorectal cancer (CRC). Nevertheless, very few reports of the association between *KALRN* mutations and cancerogenesis have been published ^12,13^.

In this study, we investigated the associations between *KALRN* mutations and anti-tumor immune signatures (NK cells, CD8+ T cells, and immune cytolytic activity (ICA)) in 10 individual cancer types (CESC, COAD, ESCA, GBM, PRAD, READ, SARC, SKCM, STAD, and UCEC) and in their pan-cancer by analyzing the Cancer Genome Atlas (TCGA, https://cancergenome.nih.gov) cohorts. In these cohorts, we also explored the associations between *KALRN* mutations and cancer immunotherapy response-related biomarkers, such as PD-L1 expression and TMB. Moreover, based on four cancer cohorts receiving immunotherapy, we investigated the association between *KALRN* mutations and cancer immunotherapy response. Furthermore, we validated our findings from bioinformatics analysis by performing in vitro experiments in three tumor cell lines (MGC803, SJSA1, and SW620). Our study demonstrated that *KALRN* mutations may promote anti-tumor immunity and cancer immunotherapy response and thus could be a useful biomarker for stratifying cancer patients responsive to immunotherapy.

## 2. Materials and Methods

### 2.1. Materials

We downloaded the datasets for the 10 TCGA cancer cohorts from the genomic data commons data portal (https://portal.gdc.cancer.gov/). The 10 cancer cohorts included cervical squamous-cell carcinoma and endocervical adeno-carcinoma (CESC), colon adeno-carcinom (COAD), Esophageal Carcinoma (ESCA), Glioblastoma Multiforme (GBM), Prostate Adenocarcinoma (PRAD), Rectum Adenocarcinoma (READ), Sarcoma (SARC), Skincutaneous Melanoma (SKCM), Stomach Adenocarcinoma (STAD), and Uterine Corpus Endometrial Carcinoma (UCEC). For each cancer type, we obtained its gene somatic mutations, RNA-Seq gene expression profiles, protein expression profiles, and clinical data. For pan-cancer, we downloaded the related datasets from UCSC (https://xenabrowser.net/datapages/). We obtained the data for predicted neoantigens in the TCGA cancer cohorts from the publication ^14^, in which a total of 6 cancer types (CESC, PRAD, GBM, STAD, SKCM, UCEC) had predicted neoantigen data available. For the four cancer cohorts receiving anti-PD-1/PD-L1/CTLA-4 immunotherapy (Allen cohort ^1^, Hugo cohort ^15^, Riaz cohort ^16^), and Rizvi cohort ^17^), we obtained their genomics and clinical data from the associated publications. A summary of these datasets is shown in Supplementary Table S1.

### 2.2. Evaluation and Comparisons of Immune Signature Enrichment Scores

We calculated the enrichment score of an immune signature in a tumor sample by the single-sample gene-set enrichment analysis (ssGSEA) ^18^ of the marker genes of the immune signature. Three immune signatures were analyzed, including NK cells (marker genes *KLRC1* and *KLRF1*), CD8+ T cells (*CD8A*), and immune cytolytic activity (*PRF1* and *GZMA*) ^14^. The comparisons of immune signature enrichment scores between two classes of samples were performed using the Mann-Whitney U test (one-sided). As well, we compared the ratios of immune-stimulatory to immune-inhibitory signatures (CD8+/CD4+ regulatory T cells, M1/M2 macrophages, and pro- /anti-inflammatory cytokines) between two classes of samples using the Mann-Whitney U test (one-sided). The ratios were the mean expression levels of immune-stimulatory signature marker genes over those of immune-inhibitory signature marker genes. We carried out these analyses using R programming software (ssGSEA scores were calculated using R package “GSVA” ^18^).

### 2.3. Comparisons of the Expression Levels of Genes and Proteins between Two Classes of Samples

We compared the expression levels of genes and proteins between two classes of samples using Student’s *t* test. The normalized RNA-Seq gene expression values (RSEM) were log2 transformed before the analyses, and the downloaded normalized protein expression data (RPPA) were used to perform the analyses.

### 2.4. Survival Analyses

We used Kaplan-Meier survival curves to exhibit the survival time differences. The significance of survival time differences was evaluated by the log-rank test. The survival analyses were performed using R function “survfit” in “survival” package.

### 2.5. In vitro Experiments

#### 2.5.1. Antibodies, Reagents, and Cell lines

All antibodies were used at a dilution of 1:1000 unless otherwise specified. Anti-PD-L1 (ab213524) was purchased from Abcam (Burlingame, CA). GAPDH (sc32233) was purchased from Santa Cruz Biotechnology. Anti-MSH2 (15520-1-AP) was purchased from Proteintech Group (Rosemont, IL, USA.). Anti-KALRN (BS3514) was purchased from Bioworld Technology (St.louis, MO). Dulbecco’s modified Eagle’s medium (12100-061), Alpha Minimum Essential Medium (MEM) (11900024), and Horse Serum (16050122) were purchased from Thermo Fisher Scientific (Waltham, MA). Inositol (I7508), 2-mercaptoethanol (M3148), and folic acid (F8758) were purchased from Sigma-Aldrich (St. Louis, MO). Human cancer cell lines MGC803 (gastric cancer), SJSA1 (osteosarcoma) and SW620 (colon cancer) were from the American Type Culture Collection (ATCC). They were cultured in a complete medium (Dulbecco’s modified Eagle’s medium supplemented with 10% fetal bovine serum) in a humidified incubator at 37°C and 5% CO_2_. NK92 cell (from KeyGEN BioTech company) was cultured in Alpha MEM with 2 mM L-glutamine, 1.5 g/L sodium bicarbonate, 0.2 mM inositol, 0.1 mM 2-mercaptoethanol, 0.02 mM folic acid, 100-200 U/ml recombinant human IL-2 (PeproTech, Rocky Hill, NJ), and a final concentration of 12.5% horse serum and 12.5% fetal bovine serum.

#### 2.5.2. Knockdown of *KALRN* with Small Interfering RNA (siRNA)

Cancer cells were transfected with *KALRN* siRNA or control siRNA using Effectene Transfection Reagent (Qiagen, B00118) following the manufacturer’s instructions. The media was replaced after 24h incubation with fresh medium and the cells were maintained for another 24h. The transfection efficiency was detected by Quantitative PCR or Western blot. *KALRN* siRNA and control siRNA were synthetized by GenePharma (Shanghai, China). Their sequences were: *KALRN* siRNA (1, 5’-GCUUCCACUGAAGUACCUATT-3’ (sense) and 5’-UAGGUAC UUCAGUGGAAGCTT-3’ (antisense); 2, 5’-GCAAUCGCCCAUUGAGUAUTT-3’ (sense) and 5’-AUACUCAAUGGGCGAUUGCTT-3’ (antisense); 3, 5’-GCUUCGACCUUGGACAC UUTT-3’ (sense) and 5’-AAGUGUCCAAGGUCGAAGCTT-3’ (antisense)), and control siRNA (5’-UUCUCCGAACGUGUCACGUTT-3’ (sense) and 5’-ACGUGACACGUUCGGA GAATT-3’ (antisense)).

#### 2.5.3. Western blotting

Cell extracts were generated using lysis buffer supplemented with protease inhibitor cocktail immediately before use. Proteins were denatured by adding 6× SDS sample buffer and boiled at 100°C for 10 min, and were then separated by SDS-PAGE. After electrophoresis, proteins were electrotransferred onto nitrocellulose membranes. The membranes were then blocked in 5% nonfat dry milk buffer at a room temperature for 1h and were then incubated overnight at 4°C with specific antibodies. Binding of the primary antibody was detected using peroxidase-conjugated secondary antibodies and a SuperSignal West Dura chemiluminescent substrate kit (Thermo Scientific, 34075) according to the manufacturer’s instructions. The western blotting result was semi-quantified using ImageJ software to measure the intensities of the bands. The band densities were normalized to background and the relative optical density ratios were calculated relative to the housekeeping gene *GAPDH*.

#### 2.5.4. Quantitative PCR

The total RNA was extracted from cells and was reversely transcribed to cDNA. Quantitative PCR was performed with the BioRad CFX96 ouchTM Real-Time PCR Detection System (BioRad, CA) using iQTM SYBR1 Green supermix (BioRad, CA). The threshold cycle numbers were obtained using BioRad CFX manager software. The program for amplification was 1 cycle of 95°C for 2 min followed by 40 cycles of 95°C for 10s, 60°C for 30s, and 72°C for 30s. The relative amount of each gene was normalized to the amount of *GAPDH*. The primer sequences were:*hKALRN*, 5’-GATTGTCATCTTCAGTGA-3’(forward) and 5’-AGGACCAAGTAATTC ATC-3’ (reverse); *hGAPDH*, 5’-TGTGGGCATCAATGGATTTGG-3’ (forward) and 5’-ACAC CATGTATTCCGGGTCAAT-3’ (reverse).

#### 2.5.5. Co-culture of Tumor Cells with NK92 Cells

The transwell chamber (Corning Inc, NY, USA) was inserted into a 6-well plate to construct a co-culture system. Tumor cells (MGC803, SJSA1, and SW620) were seeded on the 6-well plate at a density of 5×10^4^ cells/well, and NK92 cells were seeded on the membrane (polyethylene terephthalate, pore size, 0.4 μm) of the transwell chamber at a density of 5×10^4^ cells/chamber. Tumor cells and NK92 cells were co-cultured in a humidified incubator at 37°C and a 5% CO_2_ atmosphere for 48h.

#### 2.5.6. EdU Proliferation Assay

After the co-culture of tumor cells with NK92 cells for 48h, the proliferation capacity of NK92 cells was measured by an EdU (5-ethynyl-2 ‘-deoxyuridine, Invitrogen, CA, USA) proliferation assay. NK92 cells were plated in 96-well plates with a density of 2×10^3^ cells/well for 24h. Before fixation, permeabilization, and EdU staining, the cells were incubated with 10 μM EdU at 37 °C for 24h. The cell nuclei were stained with DAPI at a concentration of 1 μg/ml for 20 min. The proportion of the NK92 cells incorporating EdU was determined with fluorescence microscopy. Each assay was performed in triplicate wells.

## 3. Results

### 3.1. *KALRN* Mutations Promote Anti-Tumor Immunity in Cancer

We found that the immune signatures (NK cells, CD8+ T cells, and ICA) consistently displayed higher enrichment levels in *KALRN*-mutated than in *KALRN*-wildtype cancers in the pan-cancer analysis (Mann-Whitney U test, *P*<0.001) (Figure 1A). Moreover, the NK cells, CD8+ T cells, and ICA enrichment scores were significantly higher in *KALRN*-mutated than in *KALRN*-wildtype cancers in 4, 5, and 5 individual cancer types, respectively (Mann-Whitney U test, *P*<0.05) (Figure 1A). Furthermore, we compared the ratios of immune-stimulatory signatures to immune-inhibitory signatures in pan-cancer and in individual cancer types. We found that the ratios (CD8+/CD4+ regulatory T cells, M1/M2 macrophages, and pro-/anti-inflammatory cytokines) were remarkably higher in *KALRN*-mutated than in *KALRN*-wildtype cancers in pan-cancer and in multiple individual cancer types (Mann-Whitney U test, *P*<0.05) (Figure 1B). The EdU proliferation assay showed that the NK cells co-cultured with *KALRN*-knockdown tumor cells had significantly stronger proliferation capacity compared to the NK cells co-cultured with *KALRN*-wildtype tumor cells (Figure 1C). Taken together, these data suggest that *KALRN* mutations enhance anti-tumor immunity in cancer.

**Figure 1.**
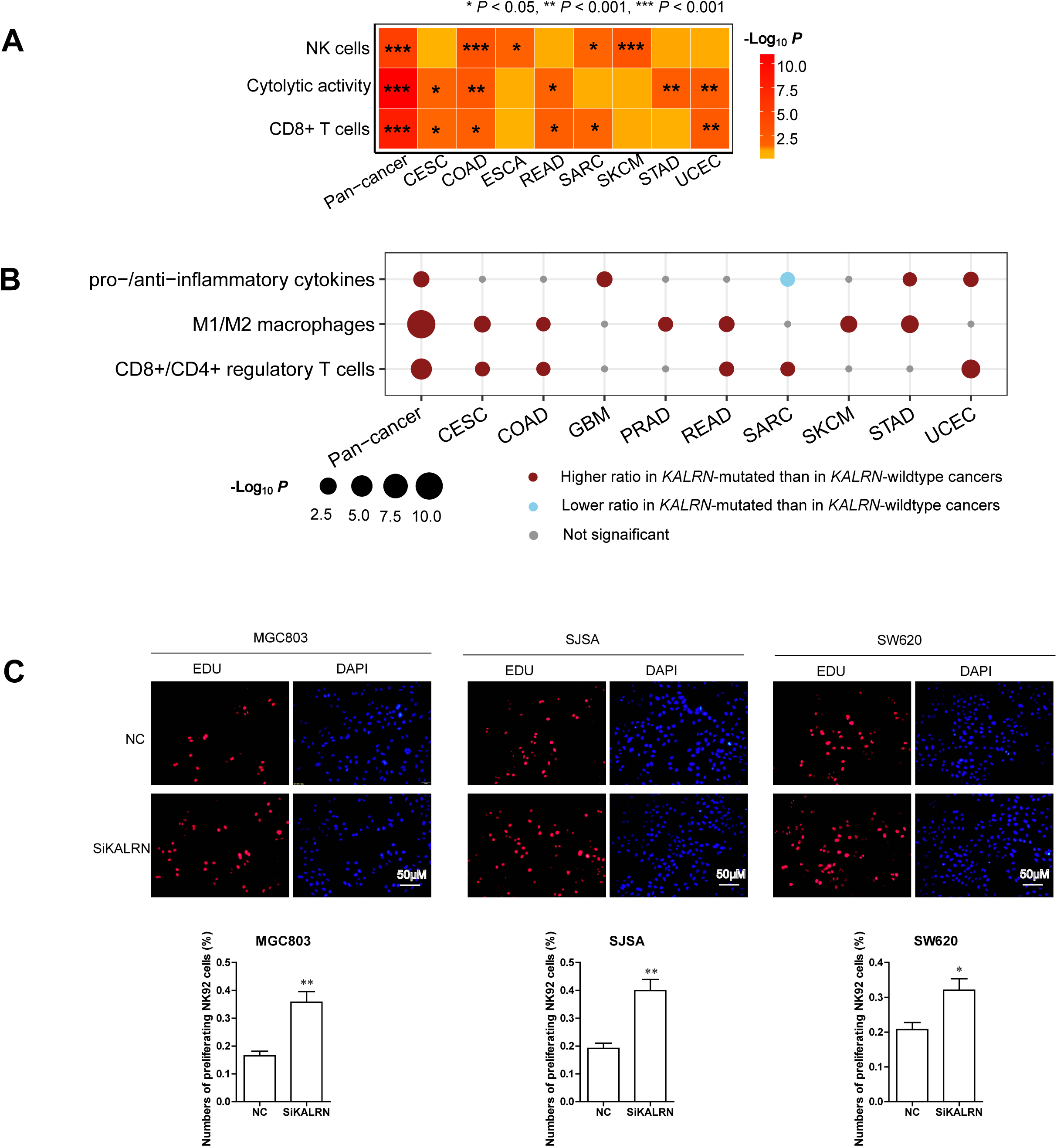
*KALRN* mutations promote anti-tumor immunity in cancer. (**A**) Three immune signatures (NK cells, CD8+ T cells, and immune cytolytic activity) show higher enrichment levels in *KALRN*-mutated than in *KALRN*-wildtype cancers in pan-cancer and in multiple individual cancer types (Mann-Whitney U test, one-sided *P*<0.05). The enrichment levels of immune signatures were evaluated by the single-sample gene-set enrichment analysis (ssGSEA) ^18^ of their marker genes. (**B**) The ratios of immune-stimulatory/immune-inhibitory signatures are significantly higher in *KALRN*-mutated than in *KALRN*-wildtype cancers in pan-cancer and in multiple individual cancer types (Mann-Whitney U test, one-sided *P*<0.05). The ratios are the mean expression levels of immune-stimulatory signature maker genes divided by the mean expression levels of immune-inhibitory signature maker genes. (**C**) The NK cells co-cultured with *KALRN*-knockdown tumor cells display significantly stronger proliferation capacity compared to the NK cells co-cultured with *KALRN*-wildtype tumor cells, evident by EdU proliferation assay. Three human cancer cell lines MGC803 (gastric cancer), SJSA1 (osteosarcoma), and SW620 (colon cancer) were used in the in vitro experiments. * *P*<0.05, ** *P*<0.01, *** *P*<0.001. It also applies to following figures.

### 3.2. *KALRN* Mutations Are Associated with Increased TMB and the Microsatellite Instability (MSI) or DNA Damage Repair Deficiency Genomic Properties in Cancer

TMB has been recognized as a predictive biomarker for cancer immunotherapy response ^19^. We found that *KALRN-*mutated cancers exhibited significantly higher TMB (defined as the total count of gene somatic mutations) than *KALRN*-wildtype cancers in pan-cancer and in all 10 individual cancer types (Mann-Whitney U test, *P*<0.01) (Figure 2A). In addition, *KALRN-*mutated cancers displayed more predicted neoantigens ^14^ than *KALRN*-wildtype cancers in pan-cancer and in 5 individual cancer types (Mann-Whitney U test, *P*<0.01) (Figure 2A).

**Figure 2.**
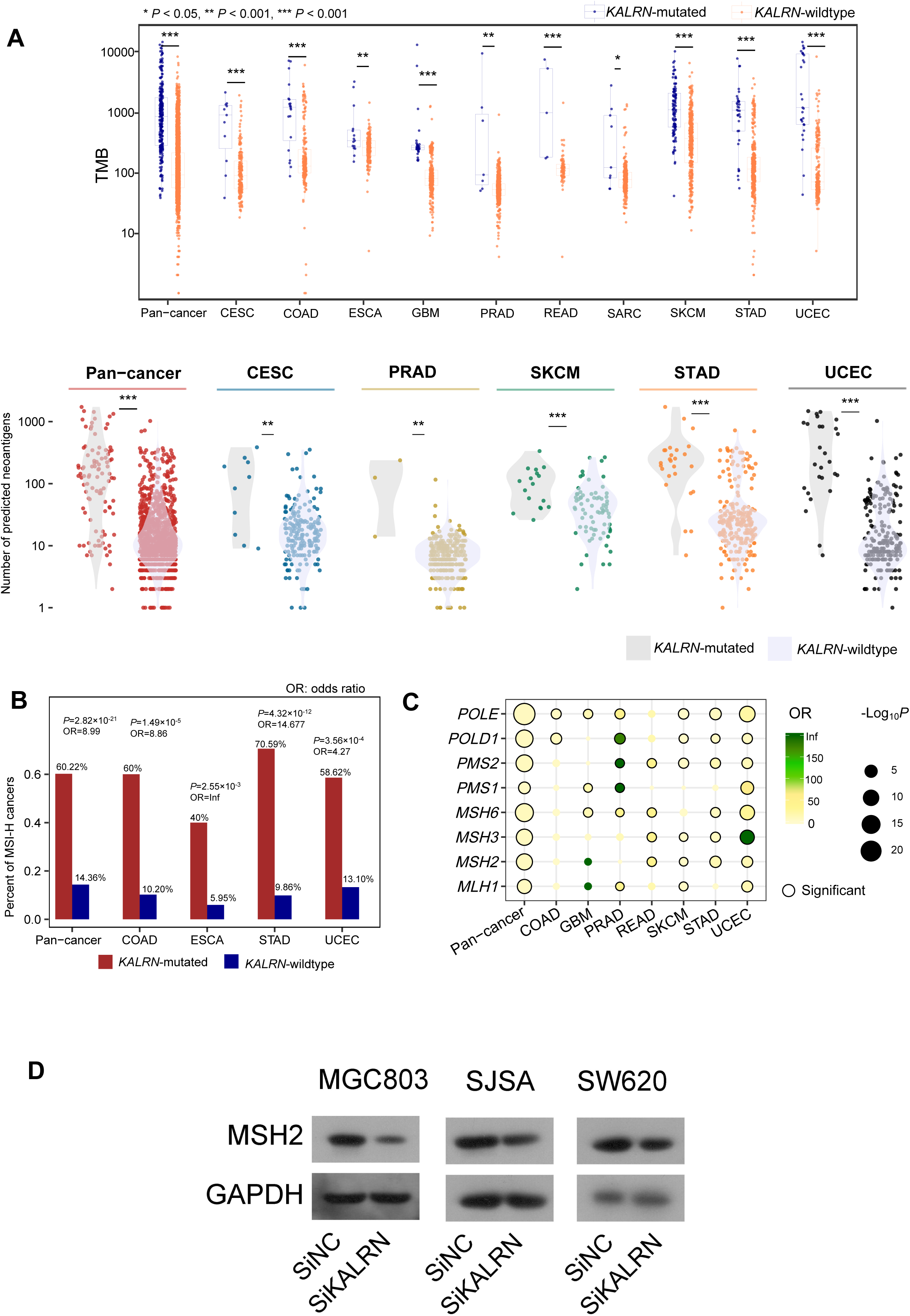
Associations of *KALRN* mutations with tumor mutation burden (TMB), neoantigens, and the microsatellite instability or DNA damage repair deficiency genomic properties in cancer. **(A)** *KALRN-*mutated cancers have significantly higher TMB and neoantigen load than *KALRN*-wildtype cancers in pan-cancer and in multiple individual cancer types (Mann-Whitney U test, one-sided *P*<0.05). TMB is defined as the total count of gene somatic mutations and the neoantigens were predicted in ^14^. **(B)** MSI-high cancers harbor more frequent mutations of *KALRN* than MSI-low/MSS cancers in pan-cancer and in 4 individual cancer types with a prevalent MSI subtype (Fisher’s exact test, *P*<0.01). MSI: microsatellite instability, MSS: microsatellite stable. They also apply to following figures. **(C)** *KALRN* mutations likely co-occur with mutations of DNA damage repair pathway genes (Fisher’s exact test, *P*<0.05). **(D)** The *KALRN*-knockdown reduced MSH2 expression in tumor cells, evident by western blotting.

Among the 5 cancer types (COAD, ESCA, READ, STAD, and UCEC) with a prevalent MSI subtype, MSI-high (MSI-H) cancers had significantly higher mutation rates of *KALRN* than MSI-low (MSI-L) and microsatellite stable (MSS) cancers in pan-cancer and in 4 individual cancer types (Fisher’s exact test, *P*<0.01) (Figure 2B). Moreover, we found that *KALRN* mutations co-occurred with mutations of DNA mismatch repair pathway genes (*PMS1, PMS2, MLH1, MSH2, MSH3*, and *MSH6*) and DNA replication and repair genes (*POLD1* and *POLE*) in pan-cancer and in multiple individual cancer types (Fisher’s exact test, *P*<0.05) (Figure 2C). By analyzing the TCGA protein expression profiling datasets with MSH2 and MSH6 expression values available, we found that MSH2 and MSH6 were downregulated in *KALRN*-mutated versus *KALRN*-wildtype cancers in UCSC and MSH6 was downregulated in *KALRN*-mutated cancers in COAD (Student’s *t* test, *P*<0.05). Furthermore, in vitro experiments showed that MSH2 expression was reduced in *KALRN*-knockdown than in *KALRN*-wildtype tumor cells (MGC803, SJSA1, and SW620) (Figure 2D). Overall, these results suggest the significant association between *KALRN* mutations and deficient DNA damage repair that could explain the significantly increased TMB in *KALRN*-mutated cancers.

### 3.3. Associations of the KALRN Downstream Target Rho Gtpases with TMB, DNA Damage Repair Pathways, and Tumor Immunity

KALRN is a member of the Rho-GEFs that activate Rho GTPases by promoting their GDP/GTP exchange ^20^. As a result, KALRN inactivation may compromise the activation of Rho GTPases. In fact, *KALRN* expression was significantly downregulated in *KALRN*-mutated versus *KALRN*-wildtype cancers in the pan-cancer analysis (Student’s *t* test, *P*=1.11×10^−5^), suggesting that *KALRN* mutations are inactivating mutations that impair KALRN activation. Network analysis by STRING ^21^ showed that KALRN interacted with multiple members of Rho GTPases, including RAC1, RAC2, RAC3, RHOA, RHOB, RHOC, RHOD, RHOG, and CDC42 (Figure 3A). Moreover, we found that *KALRN* mutations likely co-occurred with mutations of these Rho GTPase genes in pan-cancer and in 8 individual cancer types (Fisher’s exact test, *P*<0.05) (Figure 3B). Previous studies demonstrated that Rho GTPases may play a role in regulating the DNA damage response ^19^. As expected, Rho GTPase gene-mutated cancers had significantly higher TMB than Rho GTPase gene-wildtype cancers in pan-cancer and in 8 individual cancer types (Mann-Whitney U test, *P*<0.01) (Figure 3C). Rho GTPase gene*-*mutated cancers exhibited more predicted neoantigens ^14^ than Rho GTPase gene-wildtype cancers in pan-cancer and in 5 individual cancer types (Mann-Whitney U test, *P*<0.05) (Figure 3C). The mutations of Rho GTPase genes co-occurred with the mutations of DNA damage repair pathway genes (*PMS1, PMS2, MLH1, MSH2, MSH3, MSH6, POLD1*, and *POLE*) in pan-cancer and in multiple individual cancer types (Fisher’s exact test, *P*<0.05) (Figure 3D). Moreover, the mutations of Rho GTPase genes more frequently occurred in MSI-H than in MSI-L/MSS cancers in pan-cancer and in 2 individual cancer types (STAD and COAD) (Fisher’s exact test, *P*<0.05) (Figure 3E). Collectively, these results confirmed the important role of Rho GTPases in mediating DNA damage repair pathways.

**Figure 3.**
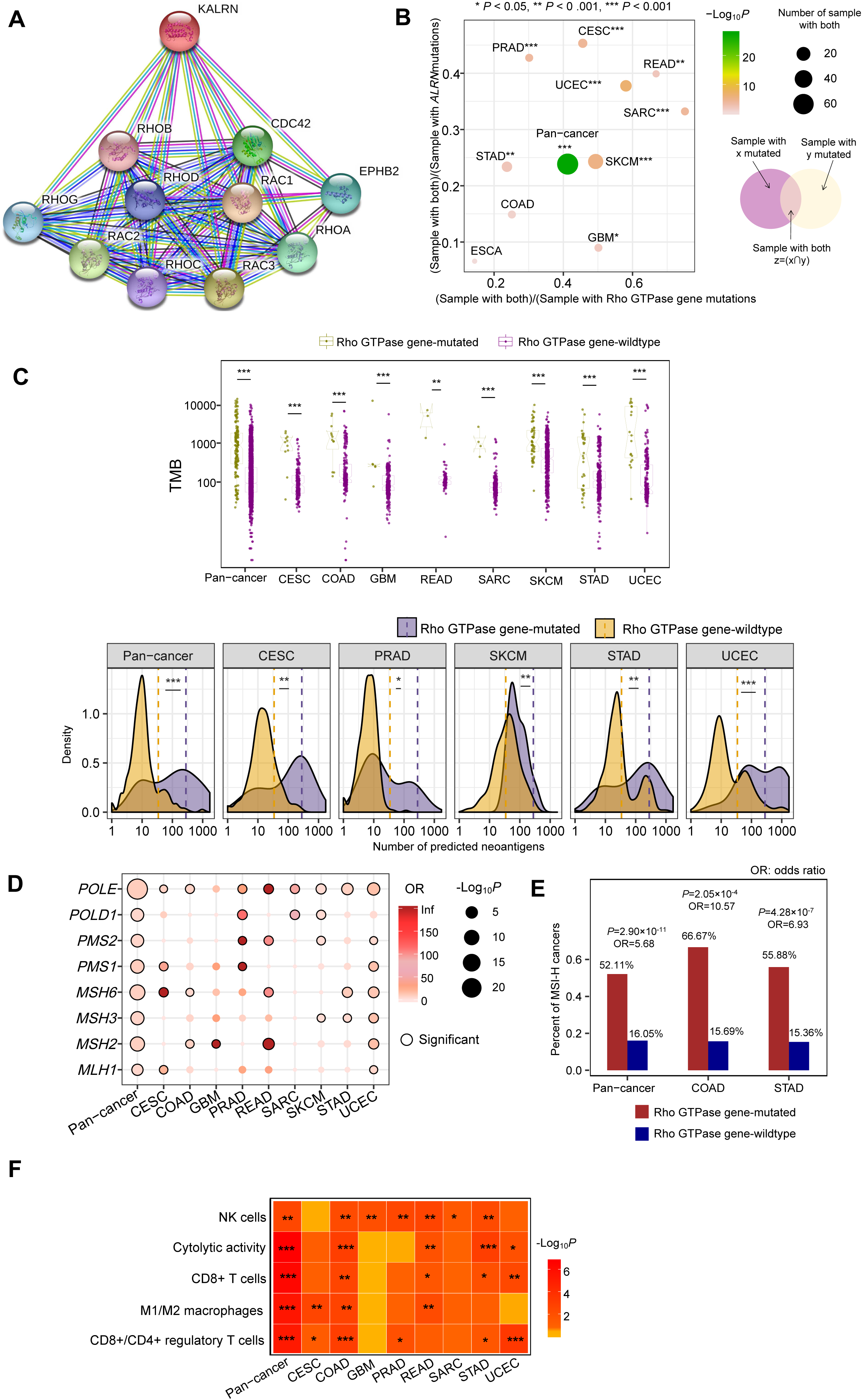
Associations of Rho GTPases with TMB, DNA damage repair pathways, and tumor immunity. **(A)** KALRN interacts with multiple members of Rho GTPases by network analysis using STRING ^21^. **(B)** *KALRN* mutations likely co-occur with mutations of Rho GTPase genes in pan-cancer and in multiple individual cancer types (Fisher’s exact test, *P*<0.05). **(C)** Rho GTPase gene-mutated cancers have higher tumor mutation burden (TMB) and neoantigen load than Rho GTPase gene-wildtype cancers in pan-cancer and in multiple individual cancer types (Mann-Whitney U test, one-sided *P*<0.01). **(D)** The mutations of Rho GTPase genes likely co-occur with the mutations of DNA damage repair pathway genes in pan-cancer and in multiple individual cancer types (Fisher’s exact test, *P*<0.05). **(E)** The mutations of Rho GTPase genes more frequently occur in MSI-high than in MSI-low/MSS cancers in pan-cancer and in 2 individual cancer types (Fisher’s exact test, *P*<0.05). **(F)** Rho GTPase gene-mutated cancers are with stronger anti-tumor immune signatures than Rho GTPase gene*-*wildtype cancers in pan-cancer and in diverse individual cancer types (Mann-Whitney U test, one-sided *P*<0.05).

Because Rho GTPase deficiency may cause DNA damage repair deficiency that in turn results in increased TMB and neoantigens, we expected that Rho GTPase gene*-*mutated cancers would display enhanced anti-tumor immune signatures. As expected, the three immune signatures (NK cells, CD8+ T cells, and ICA) were consistently stronger in Rho GTPase gene*-*mutated than in Rho GTPase gene*-*wildtype cancers in pan-cancer (Mann-Whitney U test, *P*<0.01), as well as in diverse individual cancer types (Mann-Whitney U test, *P*<0.05) (Figure 3F). Moreover, Rho GTPase gene-mutated cancers had significantly higher ratios of immune-stimulatory signatures to immune-inhibitory signatures (CD8+/CD4+ regulatory T cells, and M1/M2 macrophages) in pan-cancer and in diverse individual cancer types (Mann-Whitney U test, *P*⩽0.05) (Figure 3F).

Together, these data suggest that the significant correlations of *KALRN* mutations with increased TMB/neoantigens, deficient DNA damage repair pathways, and enhanced anti-tumor immune infiltrates could be consequences of the deregulation of Rho GTPases targeted by KALRN.

### 3.4. *KALRN* Mutations are Associated with Elevated PD-L1 Expression in Cancer and Favorable Cancer Immunotherapy Response

We found that *PD-L1* showed significantly higher expression levels in *KALRN-*mutated than in *KALRN-*wildtype cancers in pan-cancer and in 4 individual cancer types (Student’s *t* test, *P*<0.05) (Figure 4A). Furthermore, in vitro experiments showed that PD-L1 expression was significantly upregulated in *KALRN*-knockdown versus *KALRN*-wildtype tumor cells (MGC803, SJSA1, and SW620) (Figure 4B). These data suggest that PD-1–PD-L1-directed immunotherapy is likely to be more effective against *KALRN-*mutated cancers, since PD-L1 expression is a biomarker for the active response to anti-PD-1/PD-L1 immunotherapy ^6^. To prove this hypothesis, we investigated the association between *KALRN* mutations and ICB therapy response in four cancer cohorts receiving anti-PD-1/PD-L1/CTLA-4 immunotherapy, including Allen cohort ^1^ (melanoma), Hugo cohort ^22^ (melanoma), Riaz cohort ^16^ (melanoma), and Rizvi cohort ^17^ (lung cancer). We found that *KALRN*-mutated cancers had significantly higher response rates than *KALRN*-wildtype cancers in these cohorts (37.04% versus 10.96% in Allen cohort, 45% versus 11.76% in Hugo cohort, 80% versus 27.5% in Riaz cohort, and 100% versus 40.74% in Rizvi cohort) (Fisher’s exact test, *P*<0.1) (Figure 4C). Because of the appreciably higher response rates, *KALRN*-mutated cancers displayed a more favorable overall or progress-free survival tendency than *KALRN*-mutated cancers (log-rank test, *P*=0.262, 0.067, 0.055, and 0.176 for Allen, Hugo, Riaz, and Rizvi cohorts, respectively) (Figure 4D). However, the correlation between *KALRN* mutations and overall survival was much less significant in TCGA melanoma or lung cancer cohorts not receiving immunotherapy (Figure 4E). These results indicate that the more favorable ICB therapy response in *KALRN*-mutated cancer cohorts is attributed to the elevated anti-tumor immune infiltrates and PD-L1 expression in this subtype, suggesting that the *KALRN* mutation is a predictive biomarker for the active response to ICB therapy.

**Figure 4.**
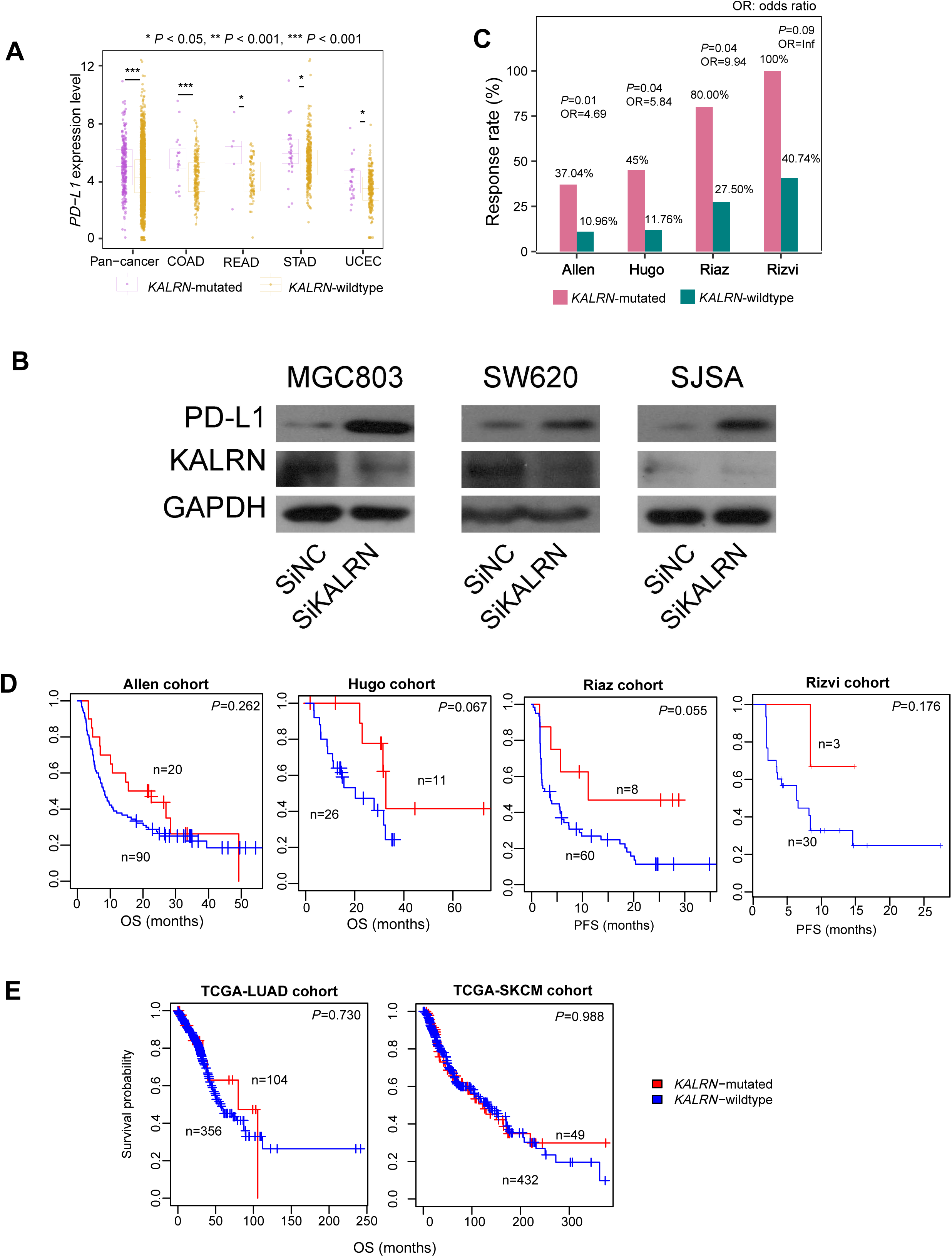
Associations of *KALRN* mutations with cancer immunotherapy response. **(A)** *KALRN-*mutated cancers display higher *PD-L1* expression levels than *KALRN-*wildtype cancers in pan-cancer and in multiple individual cancer types (Student’s *t* test, *P*<0.05). **(B)** The *KALRN*-knockdown increased PD-L1 expression in tumor cells, evident by western blotting. **(C)** *KALRN*-mutated cancers have higher response rates than *KALRN*-wildtype cancers in four cancer cohorts (Allen cohort ^1^ (melanoma), Hugo cohort ^15^ (melanoma), Riaz cohort ^16^ (melanoma), and Rizvi cohort ^17^ (lung cancer)) receiving anti-PD-1/PD-L1/CTLA-4 immunotherapy. The Fisher’s exact test *P* values and odds ratio (OR) are shown. **(D)** *KALRN*-mutated cancers show a better overall survival (OS) or progress-free survival (PFS) tendency than *KALRN*-mutated cancers in the four cancer cohorts receiving anti-PD-1/PD-L1/CTLA-4 immunotherapy. **(E)** The association between *KALRN* mutations and OS is less significant in TCGA melanoma and lung cancer cohorts without immunotherapy. In **(D)** and **(E)**, Kaplan-Meier survival curves are used to exhibit the survival time differences and the log-rank test *P* values are shown.

## 4. Discussion

For the first time, we reported the significant role of *KALRN* mutations in promoting anti-tumor immunity and immunotherapeutic vulnerabilities in diverse cancer types. The reason why *KALRN* mutations can enhance anti-tumor immunity lies in that *KALRN* mutations increase tumor mutation load to create more neoantigens due to DNA damage repair deficiency. *KALRN*-mutated cancers more highly express PD-L1, in addition to the increased anti-tumor immune infiltrates. Because both tumor-infiltrating lymphocyte density and PD-L1 expression are important factors in determining ICB therapy responsiveness ^6,23^, it is reasonable for *KALRN*-mutated cancers to exhibit increased susceptibility to ICB therapy. This was evidenced in four cancer cohorts receiving ICB therapy (Figure 4C&4D).

Furthermore, we investigated the underlying mechanism for *KALRN* mutations in enhancing anti-tumor immunity and immunotherapy response (Figure 5). The significant association between *KALRN* mutations and heightened anti-tumor immunity may be attributed to that KALRN inactivation compromise the function of KALRN target Rho GTPases on regulating DNA damage repair pathways. The function of Rho GTPases on modulating DNA damage response has been revealed in previous studies ^19^. In this study, we further demonstrated the important correlation between Rho GTPases and DNA damage repair pathways (Figure 3C, 3D&3E). Rho GTPases are involved in a wide variety of cellular functions and oncogenic processes, including cell cycle, cell death, cell migration, cell polarity, and cell adhesion ^22^,24. Notably, previous studies have shown that inhibition of Rho GTPases could promote anti-tumor immune response by increasing the immune cytolytic activity of cytotoxic CD8+ T lymphocytes ^25^. This is in accordance with our results showing that anti-tumor immune signatures grew stronger in the cancers harboring mutations of Rho GTPase genes (Figure 3F). Interestingly, we found that the cancers with mutations of Rho GTPase genes over-expressed *PD-L1* in pan-cancer and multiple individual cancer types (Mann-Whitney U test, *P*<0.05). It suggests that mutations of Rho GTPase genes may favor ICB therapy response. Indeed, we found that the cancers harboring mutations of Rho GTPase genes showed higher ICB therapy response rates than those without such mutations in the four cancer cohorts receiving ICB therapy (42.86% versus 22.78% in Allen cohort, 70% versus 48.15% in Hugo cohort, 83.33% versus 25.64% in Riaz cohort, and 100% versus 45% in Rizvi cohort).

**Figure 5.**
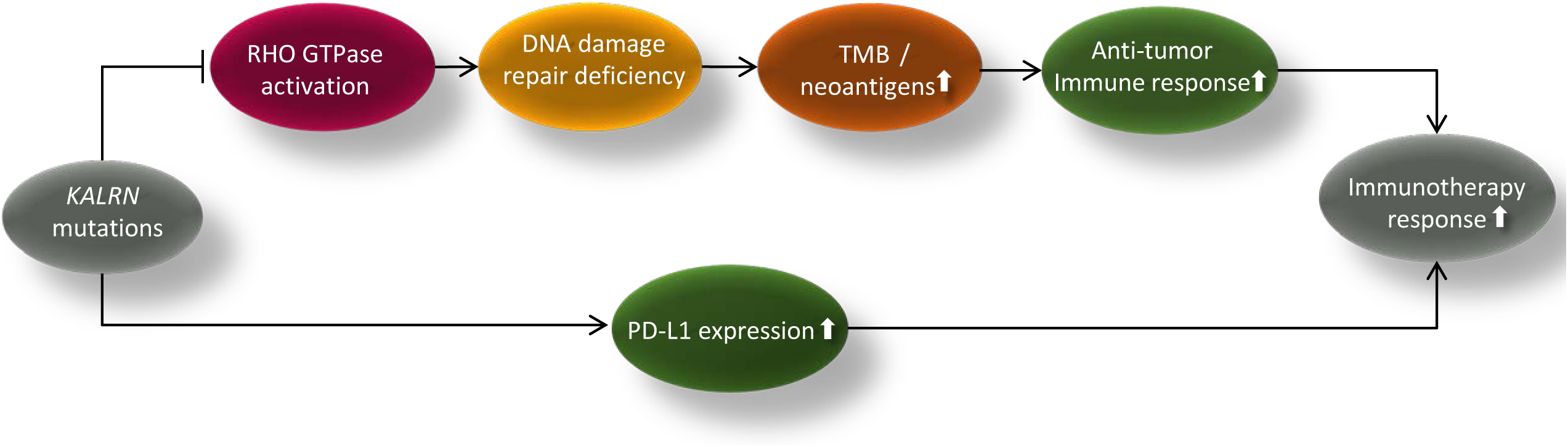
The mechanism that *KALRN* mutations promote anti-tumor immunity and immunotherapy response in cancer.

## 5. Conclusions

The *KALRN* mutation correlates with enhanced anti-tumor immunity and immunotherapy response and thus is a predictive biomarker for the response to immunotherapy.

## List of abbreviations

*KALRN*: Kalirin RhoGEF kinase;
NK: Natural Killer;
ICA: Immune Cytolytic Activity;
ICB: Immune Checkpoint Blockade;
CESC: Cervical Squamous-Cell Carcinoma And Endocervical Adeno-Carcinoma;
CRC: Colorectal Cancer;
COAD: Colon Adenocarcinoma;
ESCA: Esophageal Carcinoma;
GBM: Glioblastoma Multiforme;
PRAD: Prostate Adenocarcinoma;
READ: Rectum Adenocarcinoma;
SARC: Sarcoma;
SKCM: Skincutaneous Melanoma;
STAD: Stomach Adenocarcinoma;
UCEC: Uterine Corpus Endometrial Carcinoma;
MSI: Microsatellite Instability;
MSS: Microsatellite stable;
OR: Odds Ratio;
OS: Overall Survival;
PFS: Progress-Free Survival;
CTLA4: Cytotoxic T-lymphocyte-associated protein 4;
PD-1: Programmed cell death protein 1;
PD-L1: Programmed cell death 1 ligand;
ssGSEA: single-sample Gene-Set Enrichment Analysis;
RSEM: RNA-Seq gene expression values;
RPPA: Reverse Phase Protein Array;
TCGA: The Cancer Genome Atlas;
TMB: Tumor Mutation Burden;
ATCC: American Type Culture Collection;
siRNA: Small Interfering RNA;
PCR: Polymerase chain reaction.

## Authors’ Contributions

ML performed experiments and data analyses, and helped prepare for the manuscript. YM performed experiments and data analyses, and helped prepare for the manuscript. YZ performed experiments. LQ performed experiments. XW conceived this study, designed analysis strategies, and wrote the manuscript. All the authors read and approved the final manuscript.

## Acknowledgments

We thank Ms. Yin He for help in performing the experiments.

## Supplementary Data

**Table S1.**
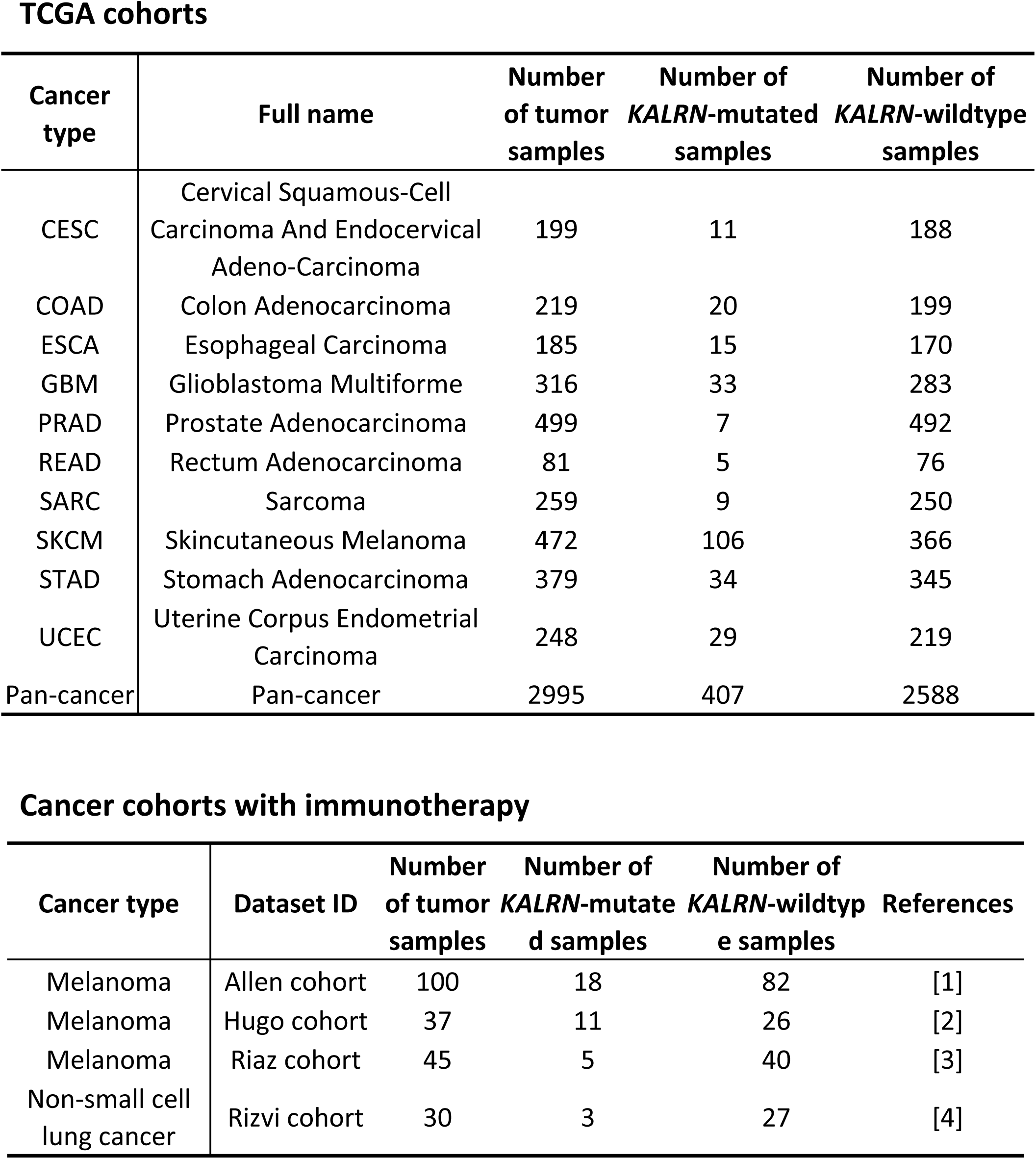
A summary of the datasets used in this study.

